# Spontaneous mutations in *braRS* and *braAB* and the IS*1181*-related deletion of *vraDE* genes increase the susceptibility of *Staphylococcus aureus* clinical isolates to epidermin

**DOI:** 10.1101/2025.09.27.678946

**Authors:** Yujin Suzuki, Airi Odawara, Miki Kawada-Matsuo, Junzo Hisatsune, Chika Arai, Saki Nishihama, Hideki Shiba, Motoyuki Sugai, Souichi Yanamoto, Takemasa Sakaguchi, Hitoshi Komatsuzawa

## Abstract

Bacteriocins, which are antimicrobial factors produced by bacteria, are expected to be candidates for a novel class of antimicrobial agents against antimicrobial-resistant bacteria, including methicillin-resistant *Staphylococcus aureus*. To evaluate its effectiveness, the examination of susceptibility using multiple clinical isolates is necessary. In this study, we focused on epidermin, a class I bacteriocin produced by *Staphylococcus epidermidis*. Among the 146 *S. aureus* clinical isolates, 77% did not show susceptibility to epidermin, but the susceptibility of some of the strains was significantly higher. Knowing that the two-component regulatory system BraRS affects the susceptibility to epidermin, we analyzed the inducibility of the effector gene *vraD* in susceptible strains and found that its ability is lost in highly susceptible strains. Some of these strains harbor frameshift or nonsense mutations in BraR, BraA and BraB. Interestingly, the *vraDE* genes with 35,005 bp neighboring genomic regions, including the *icaRADBC* and histidine biosynthesis operons, were lost in some highly susceptible strains. In addition, all the *vraDE*-deficient strains were CC121, and IS*1181* sequences located immediately upstream and downstream of the deleted region were found in some *vraDE*-positive CC121 strains. The deleted region was replaced by one copy of the IS*1181* sequence in *vraDE*-deficient strains. Subculturing of the *vraDE*-positive CC121 strain generated a *vraDE-*deficient mutant with increased susceptibility to epidermin at a rate of approximately 1%, suggesting that recombination between IS*1181* sequences results in the deletion of the genomic region, including *vraDE,* in some CC121 strains.

## Introduction

*Staphylococcus aureus* is an opportunistic pathogen that sometimes causes several infections, such as impetigo, toxic shock syndrome, endocarditis and sepsis (1, 2). In addition, methicillin-resistant *S. aureus* (MRSA) is known as a widespread antimicrobial-resistant bacterium. In most cases, MRSA carries not only the methicillin resistance gene *mecA* but also resistance genes against other classes of antimicrobial agents (3). Since the chemotherapy for MRSA infection depends on limited antimicrobial agents such as vancomycin, daptomycin and linezolid (4), novel antimicrobial agents that are effective against MRSA are expected. Bacteriocins, antimicrobial peptides or proteins produced by bacteria are among the candidates. In addition, bacteriocin-producing bacteria are thought to act against competing bacteria in their niche and contribute to securing and expanding their own survival area (5), and *Staphylococcus epidermidis,* a commensal bacterium that frequently coexists with *S. aureus,* is known to produce bacteriocins that have anti-*S. aureus* activity (6, 7). Among them, epidermin has been reported to be effective against multiple *S. aureus* strains (7, 8). Despite growing interest in bacteriocins, studies investigating the link between genomic information and bacteriocin susceptibility in the clinical isolates remain limited.

Some two-component regulatory systems (TCSs) affect the susceptibility of *S. aureus* to bacteriocin. The BraRS system, which is known to confer resistance to nisin A, nukacin ISK-1, bacitracin and epidermin (9–11), senses the presence of bacteriocins in concert with the sensor transporter BraAB and induces gene expression of the *vraDE* ABC transporter, which strongly effluxes bacteriocins (12). GraRS is another TCS that confers resistance to cationic antimicrobial agents such as polymyxin B, nisin A, nukacin ISK-1 and Pep5 by inducing the expression of *vraFG, dlt*-operon and *mprF* (9, 11, 13). VraFG is an ABC transporter that binds with GraRS to induce the expression of downstream genes(14, 15). Dlt couples D-alanine with teichoic acids, and MprF couples L-lysine with phosphatidylglycerol, thereby weakening the net surface negative charge and repelling cationic antimicrobial agents by electrostatic repulsion (16, 17). VraSR, a TCS that confers resistance to nukacin ISK-1 (9), senses obstacles in cell wall synthesis caused by cell wall inhibitors and induces the expression of cell wall synthesis-related genes such as *sgtB, pbpB* and *murZ* (9, 18). In addition, deficiency in the SrrAB system results in decreased susceptibility to Pep5, nisin A and nukacin ISK-1 but not to epidermin because of decreased aerobic respiration (11).

In this study, we examined the variation in epidermin susceptibility among 146 *S. aureus* clinical isolates and verified the underlying mechanisms of the difference in susceptibility among strains using genomic information.

## Results

### Distribution of epidermin susceptibility in *S. aureus* clinical isolates

To verify epidermin susceptibility among various *S. aureus* strains, we performed a direct assay using 146 *S. aureus* clinical isolates as indicators and the epidermin-producing *S. epidermidis* strain KSE56 as the producer strain. This collection contains strains isolated from the nasal cavity of healthy people (KSA, HSA strains) and patients with bloodstream infections (19–21). The susceptibility of the *S. aureus* strains was determined by measuring the diameter of the inhibition zone (halo). Since the diameter of the producer colony was 6 mm, no halo was defined as 6 mm. Approximately 77.4% of the strains (113 strains) presented no halo, but 33 strains presented halos (Fig. 1). Among them, the halo sizes varied greatly, and the largest halo size was 28.19 mm (HSA94).

**Fig. 1.**
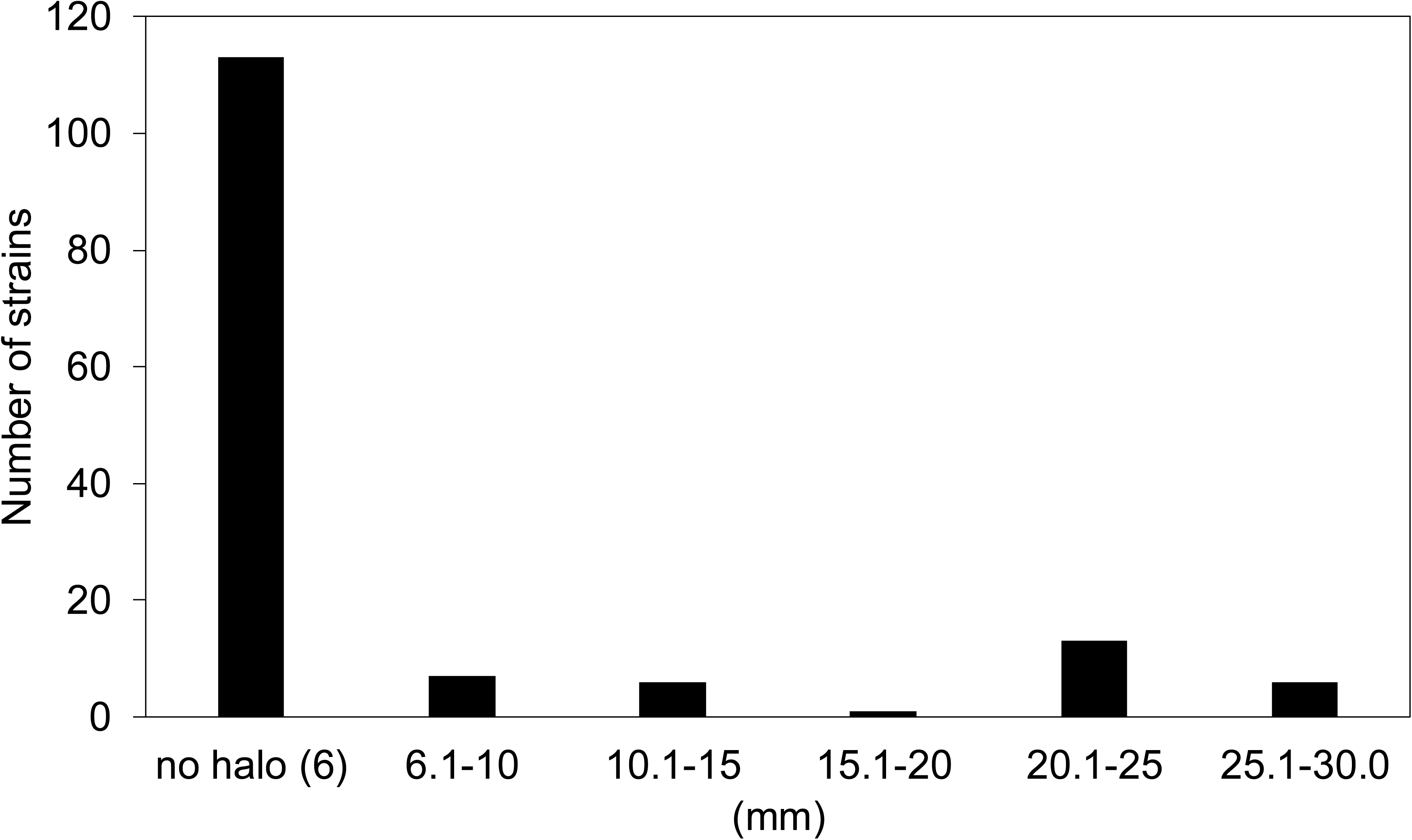
Distribution of the halo size of epidermin in 146 *S. aureus* clinical isolates. The bar graphs depict the distribution of halo sizes against KSE56. Since 6 mm is a spotted KSE56 colony size, it was determined to have no halo.

### BraRS and its regulated genes confer resistance against epidermin

Since differences in epidermin susceptibility were detected among clinical isolates, it was predicted that differences in the function of the factor affecting epidermin susceptibility caused the differences in susceptibility among strains. In *S. aureus,* several two-component regulatory systems (TCSs) affect bacteriocin susceptibility (9). Previously, we demonstrated that BraRS confers resistance against epidermin (11), but the influence of other TCSs on the susceptibility to epidermin has not been revealed. Thus, we performed a direct assay using the *braRS*-inactivated mutant (Δ*braRS*), *graRS*-inactivated mutant (Δ*graRS*) and *vraSR*-inactivated mutant (Δ*vraSR*) of the *S. aureus* MW2 strain as indicators. Compared with the wild-type (WT) strain, only the Δ*braRS* mutant showed significantly increased susceptibility to epidermin (Fig. 2). We also analyzed the susceptibility of the *braAB*-inactivated mutant (Δ*braAB*) and *vraDE*-inactivated mutant (Δ*vraDE*) with the factors regulated by and associated with BraRS and found that these mutants showed increased susceptibility to epidermin. Since BraRS is known to induce the expression of the *vraDE* gene in the presence of nisin A and nukacin ISK-1 (9, 22), we verified whether the presence of epidermin also causes the induction of *vraDE. vraD* expression was significantly induced in the presence of 128 μg/mL epidermin in the MW2 WT, as was observed in the presence of nisin A. However, *vraD* expression was not induced in the presence of nisin A or epidermin in the MW2 Δ*braRS* mutant (Fig. S1).

**Fig. 2.**
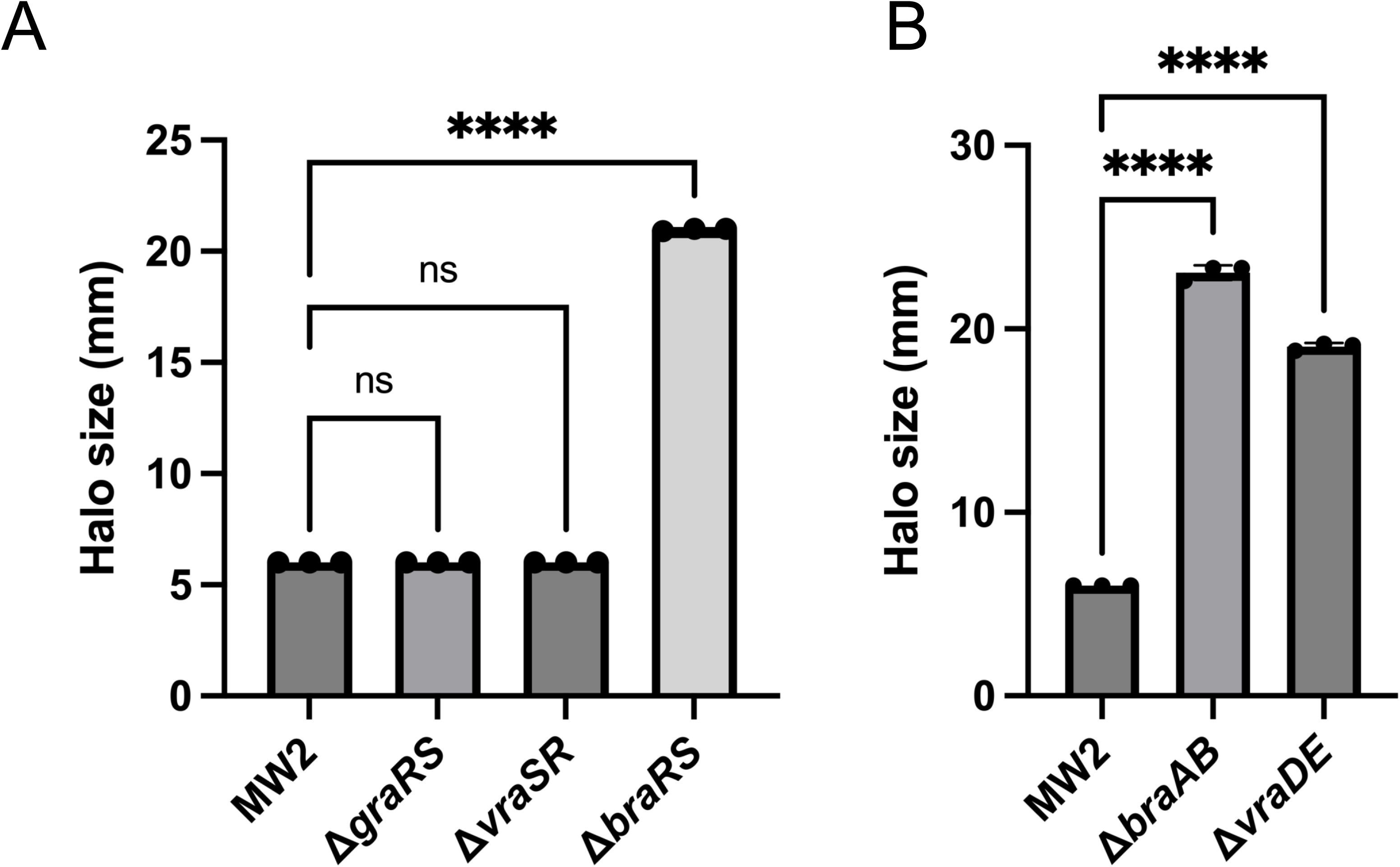
Epidermin susceptibility in MW2 gene-inactivated mutants. (A) Comparison of the halo sizes of MW2 WT and 3 TCS-inactivated mutants against *S. epidermidis* KSE56. The exact halo size was determined from the mean value of three independent experiments. Statistical significance was determined by Dunnet’s multiple comparison test. ****, *P <* 0.0001; ns, not significant. (B) Comparison of the halo sizes of the MW2 WT, Δ*braAB,* and Δ*vraDE* mutants against KSE56. The exact halo size was determined from the mean value of three independent experiments. Statistical significance was determined by Dunnet’s multiple comparison test. ****, *P <* 0.0001; ns, not significant.

### Inducibility of *vraD* was lower in epidermin-susceptible strains

Since the BraRS system strongly affects epidermin susceptibility in *S. aureus* and some clinical strains are highly susceptible, we analyzed the inducibility of *vra*D gene expression in the presence of nisin A in epidermin-susceptible strains and MW2 as a control strain (epidermin-nonsusceptible). The halo sizes of all the strains with halos and MW2 are shown in Fig. 3A. Gene expression was quantified by RT‒qPCR using *vraD*-specific primers in the presence or absence of 128 μg/mL nisin A in those strains. Fig. 3B (halo size: greater than 20 mm) and 3C (halo size: less than 20 mm, greater than 6 mm) show *vraD* expression in the presence (nis+) or absence of nisin A. In highly susceptible strains with a halo size of 20 mm or greater, there was no upregulation of *vraD* expression in the presence of nisin A, whereas an increase in expression was observed in strains with relatively small halo sizes and in MW2. In addition, *vraD* was negative based on PCR in three highly susceptible strains, HSA44, KSA152 and 33009-19-S058 (analyzed in the following section).

**Fig. 3.**
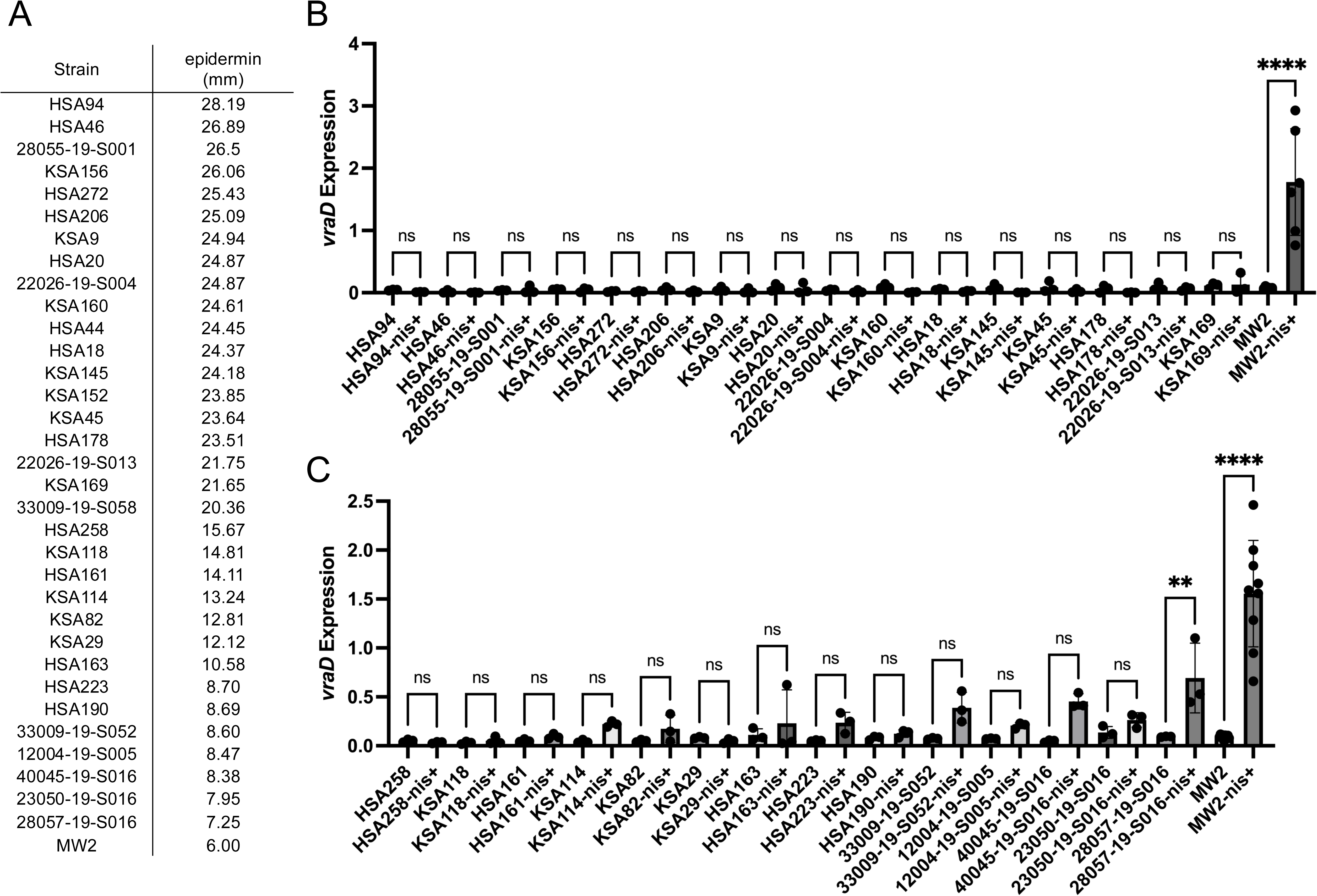
Gene expression of *vraD* with or without nisin A in *S. aureus* clinical isolates that had halos. (A) Halo sizes of all strains with halos. The strains were sorted by halo size. MW2 is shown as a control strain. (B) The gene expression of *vraD* in all strains with halo sizes greater than 20 mm in the presence (nis+) or absence of 128 µg/mL nisin A. Statistical analysis was performed using Šidák’s multiple comparison test. ****, *P* < 0.0001; ns, not significant. (C) The gene expression of *vraD* in all strains with halos smaller than 20 mm in the presence or absence of 128 µg/mL nisin A. Statistical analysis was performed using Šidák’s multiple comparison test. **, *P* < 0.01; ****, *P* < 0.0001; ns, not significant.

To elucidate the reason for the loss of *vraD* induction, we analyzed the amino acid sequences of BraR, BraS, BraA and BraB. Several amino acid substitutions were detected in most of the strains, but mutations known to affect the activity or expression of *braRS* identified in previous studies were not detected in all clinical isolates (23–25). Interestingly, in 12 highly susceptible strains, frameshifts or nonsense mutations were detected in *braR, braA* and *braB* (Fig. 4). In addition, considering the possibility that amino acid substitutions in BraR, BraS, BraA, and BraB are involved in loss of function, mutations specific to these strains were examined, and specific substitutions in HSA46, HSA272, HSA178, HSA258, KSA114, KSA29 and 40045-19-S016 were detected (Fig. 4). The positions of each substitution are plotted in Fig. 4B.

**Fig. 4.**
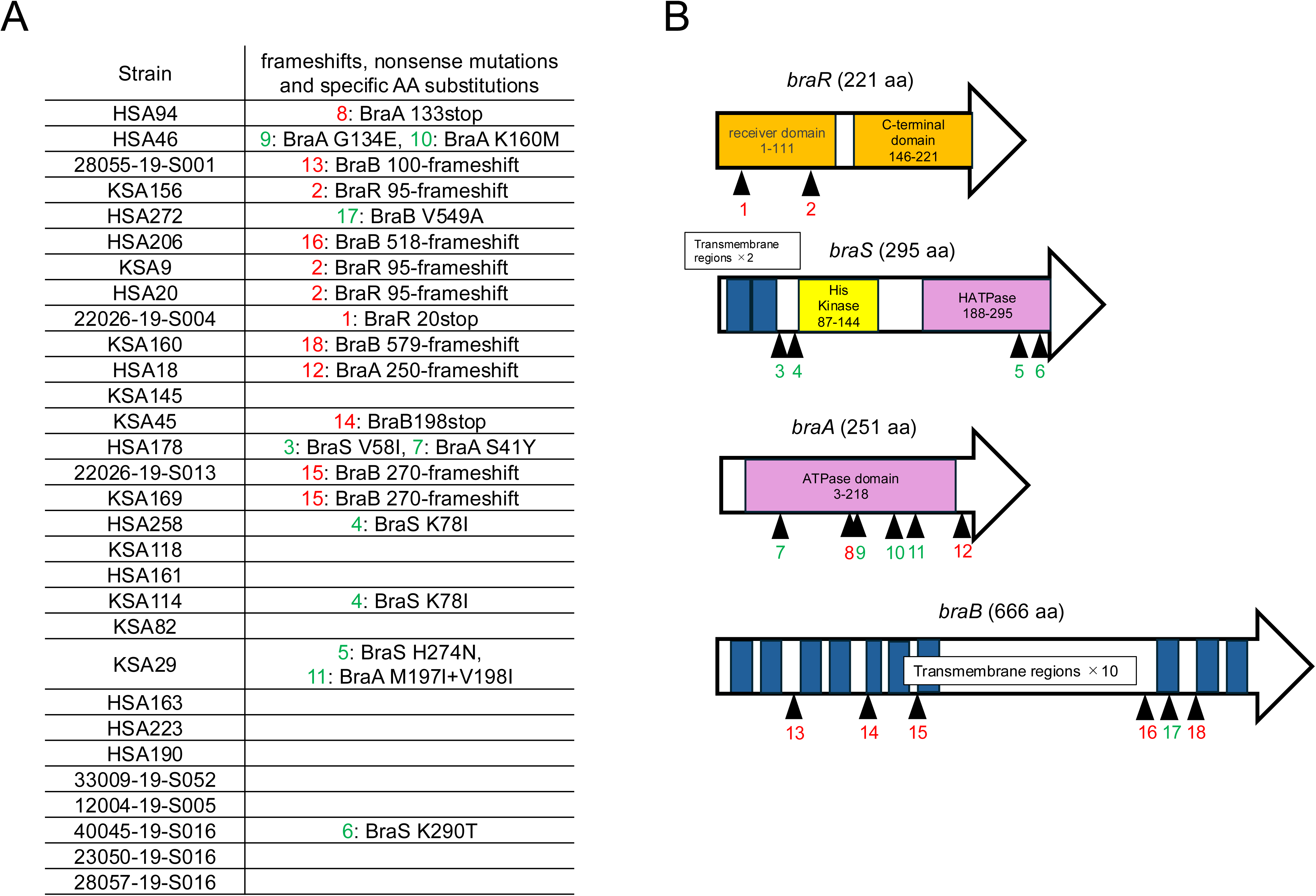
Genetic mutations in the *braRS/braAB* genes in strains with halos. (A) Positions of frameshifts, nonsense mutations (red) and specific amino acid substitutions (green) in strains with halos. The initial numbers correspond to the positions of the mutations depicted in Figure B. (B) The positions of each mutation in BraRS/BraAB. The boxes within the arrows indicate the functional domains.

### Missing *vraDE* on the chromosome in some CC121 strains

The whole-genome sequences of three strains without amplification of *vraD* by PCR (KSA152, HSA44, and 33009-19-S058) revealed no significant *vraDE* gene hits in the BLASTn analysis. All these strains were classified as clonal complex (CC) 121 (the distribution of CCs in our collection is shown in Fig. S2). To investigate the length of the missing region containing *vraDE* in these strains, we determined the complete genome sequence of three strains and the control strain (KSA13), which is the same lineage (CC121) and possessing *vraDE*. Genomic comparison revealed that the region from 393 bp upstream of the end of *icaR* to 78 bp downstream of *cspB* (35,005 bp in total) was missing in three strains (Fig. 5). The missing genomic region contained other genes, such as the biofilm formation-related genes *icaRADBC* and histidine biosynthesis gene cluster *his*-operon. Consistent with the genome data, the three strains showed histidine auxotrophy in chemically defined medium (CDM) (Fig. 6), but no difference in the strength of biofilm formation was observed among the CC121 strains (Fig. S3). Interestingly, the IS*1181* copies were present immediately upstream and downstream of the missing region in KSA13 (Fig. 5), and the missing region in the three strains was replaced by one copy of the IS*1181* sequence. The nucleotide sequences of the two copies of IS*1181* in KSA13 were completely identical (data not shown).

**Fig. 5.**
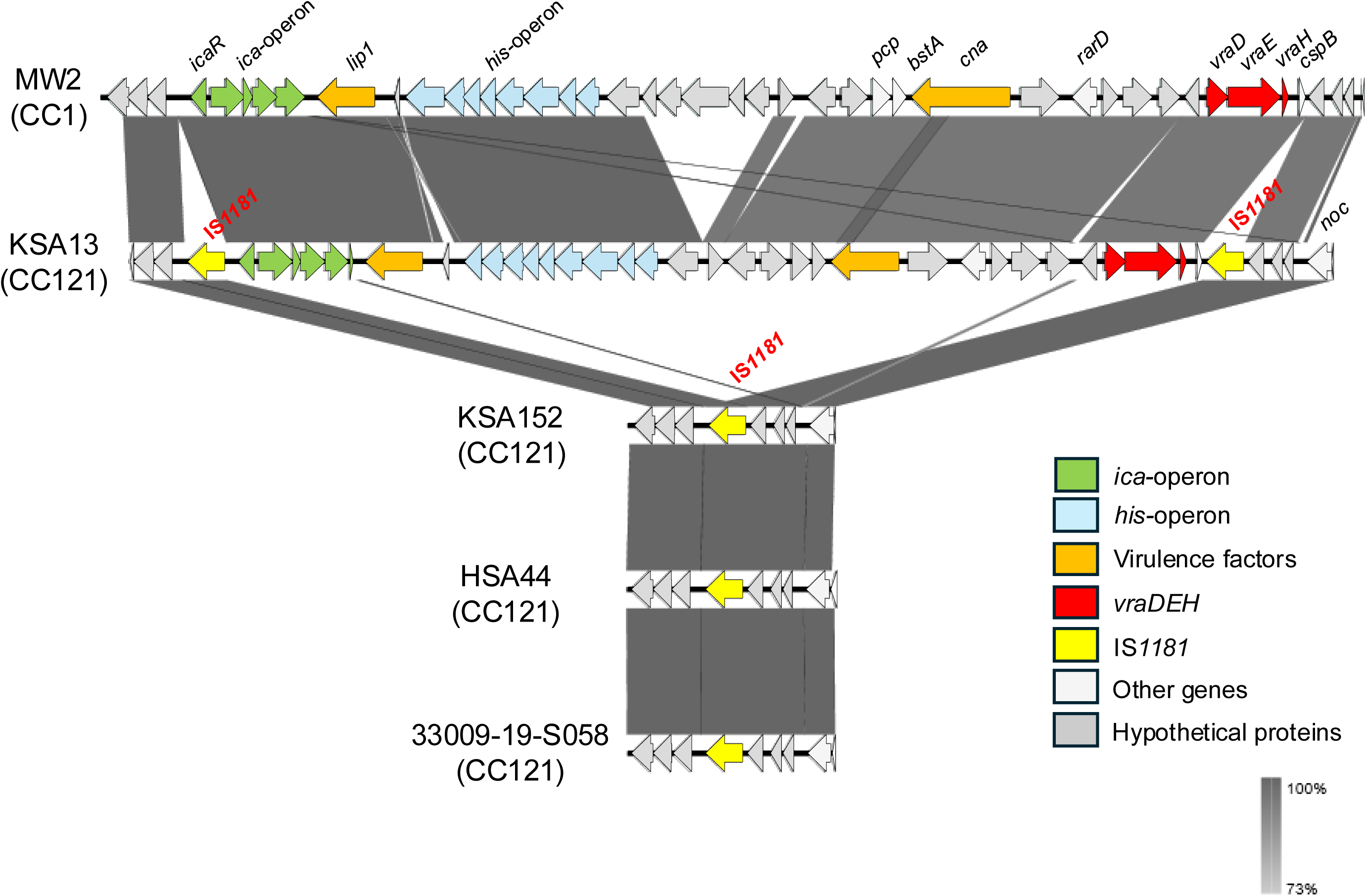
Comparison of gene arrangements adjacent to the *vraDE* gene in the *vraDE*-missing strains. Comparison of the gene arrangements of missing regions in KSA152, HSA44 and 33009-19-S058. MW2 and KSA13 are shown as *vraDE-*positive controls. The arrows represent the gene loci, and grey shading indicates the homology of DNA sequences. The legend for the arrow colors is shown on the lower right.

**Fig. 6.**
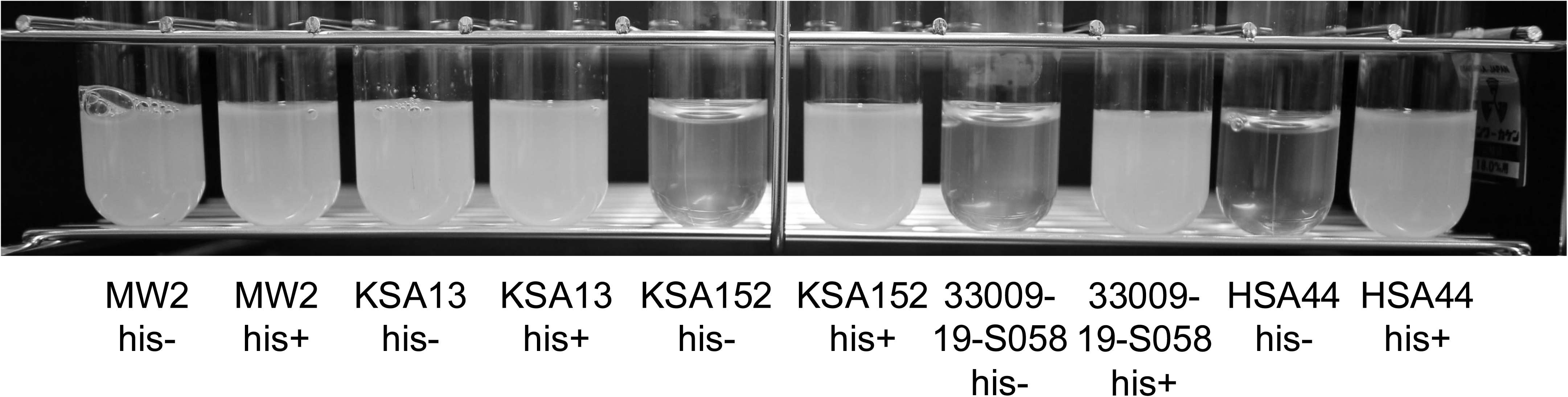
Growth of *S. aureus* strains in CDM with or without histidine. MW2, KSA13, KSA152, 33009-19-S058 and HSA44 were cultured in 3 mL of CDM with (his+) or without (his-) 1 mM L-histidine for 24 h with shaking.

### *ica, his* and *vraDE*-deleted mutants were obtained from the subculture of the CC121 strain

In KSA13, two identical IS*1181* copies sandwiched the missing region observed in KSA152, HSA44, and 33009-19-S058, and we speculated that homologous recombination could occur between these ISs and that the genomic region, including *ica, his* and *vraDE,* could be deleted. To verify this hypothesis, we cultured KSA13 for ten passages and obtained 396 colonies. Among them, three colonies exhibited histidine auxotrophy and increased susceptibility to epidermin (Fig. 7A, 7B). PCR using *vraD, icaA,* and *hisD*-specific primers resulted in no amplification of the three mutants, whereas KSA13 resulted in amplification of all the genes (data not shown). DNA sequencing of the focused region in mutant 1 revealed that this mutant had the same genomic deletion as the KSA152, HSA44, and 33009-19-S058 strains (Fig. 7C).

**Fig. 7.**
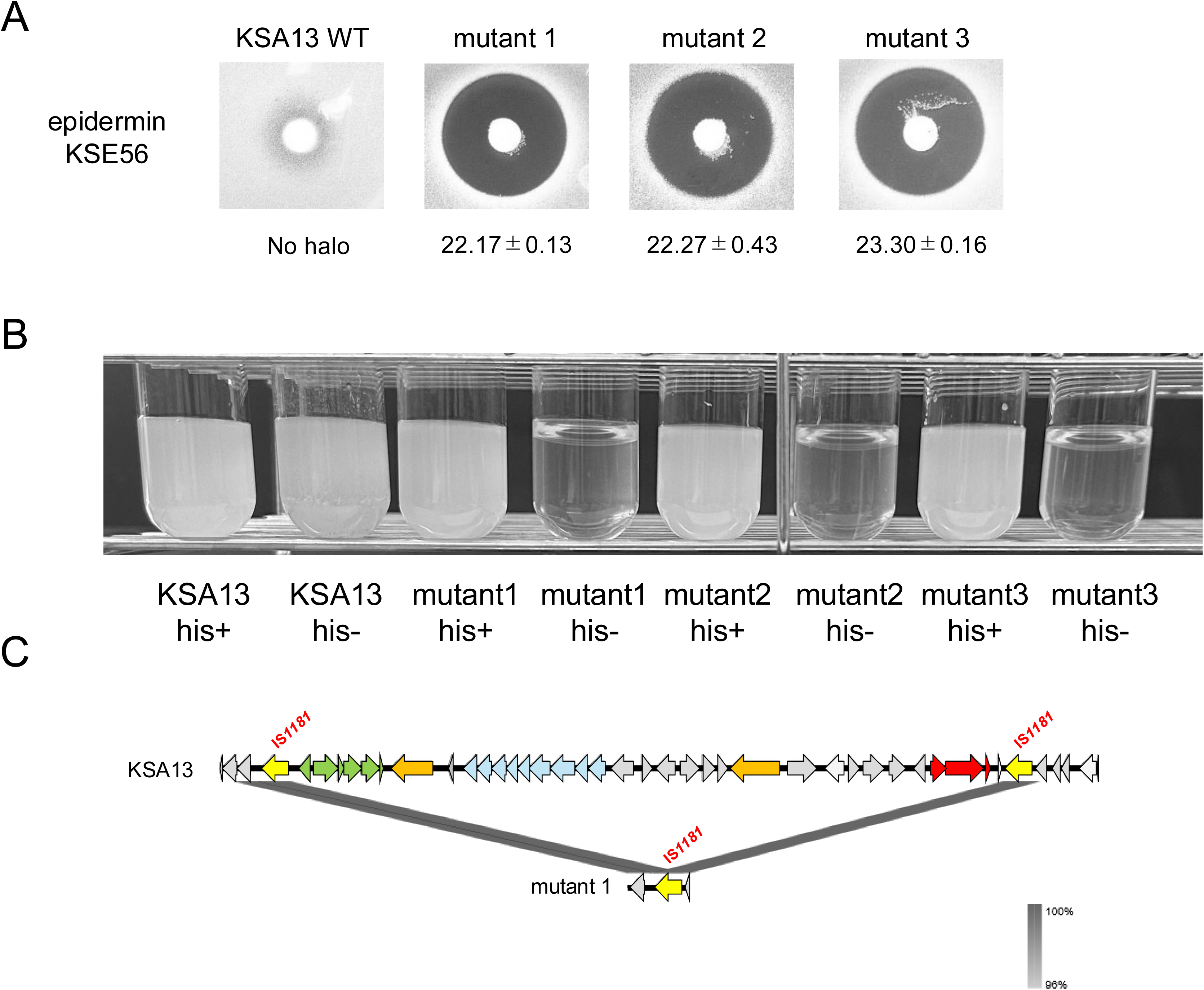
Analysis of KSA13 and its *vraDE-*deleted mutant. (A) Halos of KSA13 WT and three obtained mutants. The halo sizes and their standard deviations are shown below the figures. (B) Growth of KSA13 WT and three mutants in CDM with or without 1 mM L-histidine. The strains were cultured in 3 mL of CDM with or without 1 mM L-histidine for 24 h with shaking. (C) Comparison of the gene arrangements of WT and mutant 1 of KSA13. The arrows represent the gene loci, and grey shading indicates the homology of DNA sequences. The legend for the arrow colors is the same as that in Fig. 5.

### IS*1181* around *vraDE* was specifically found in CC121

IS*1181* is known to be present in multiple copies on the chromosome of *S. aureus* (26). Since the presence of IS*1181* at specific positions is predicted to be necessary for genomic deletion, we verified (i) the presence of the *vraDE* gene and (ii) the numbers and positions of IS*1181* on publicly available *S. aureus* genomes to investigate whether the loss of *vraDE* or the presence of IS*1181* around *vraDE* are common in *S. aureus*. Since the multiple copies of IS*1181* on a chromosome may sometimes cause a loss of contig continuity, we used publicly available genome data with a ‘complete’ assembly level for this analysis. BLASTn analysis revealed that there was no strain with missing *vraDE* in the analyzed genomes of 1406 strains. With respect to IS*1181*, the number and positions of IS*1181* copies differed among strains. Compared with the lineage strains, the CC5, CC97, and CC121 strains presented relatively high numbers of IS*1181* copies, the CC1 and CC8 strains presented relatively low numbers, and the CC15, CC22, CC30, and CC45 strains presented no copies (Fig. S4). Within the same CC, the number of IS*1181* copies varied among strains, and some of the inserted positions differed, but some of them were conserved among strains (Fig. 8). Specifically, IS*1181*, located downstream of *cspB,* was found in 14 strains, and all of them were CC121. Among them, only one strain had two IS*1181* genes located both upstream of *icaR* and downstream of *cspB* (Fig. 8) (strain SA11-SX) (27). In addition, three strains of CC121 did not have an IS*1181* around *vraDE*.

**Fig. 8.**
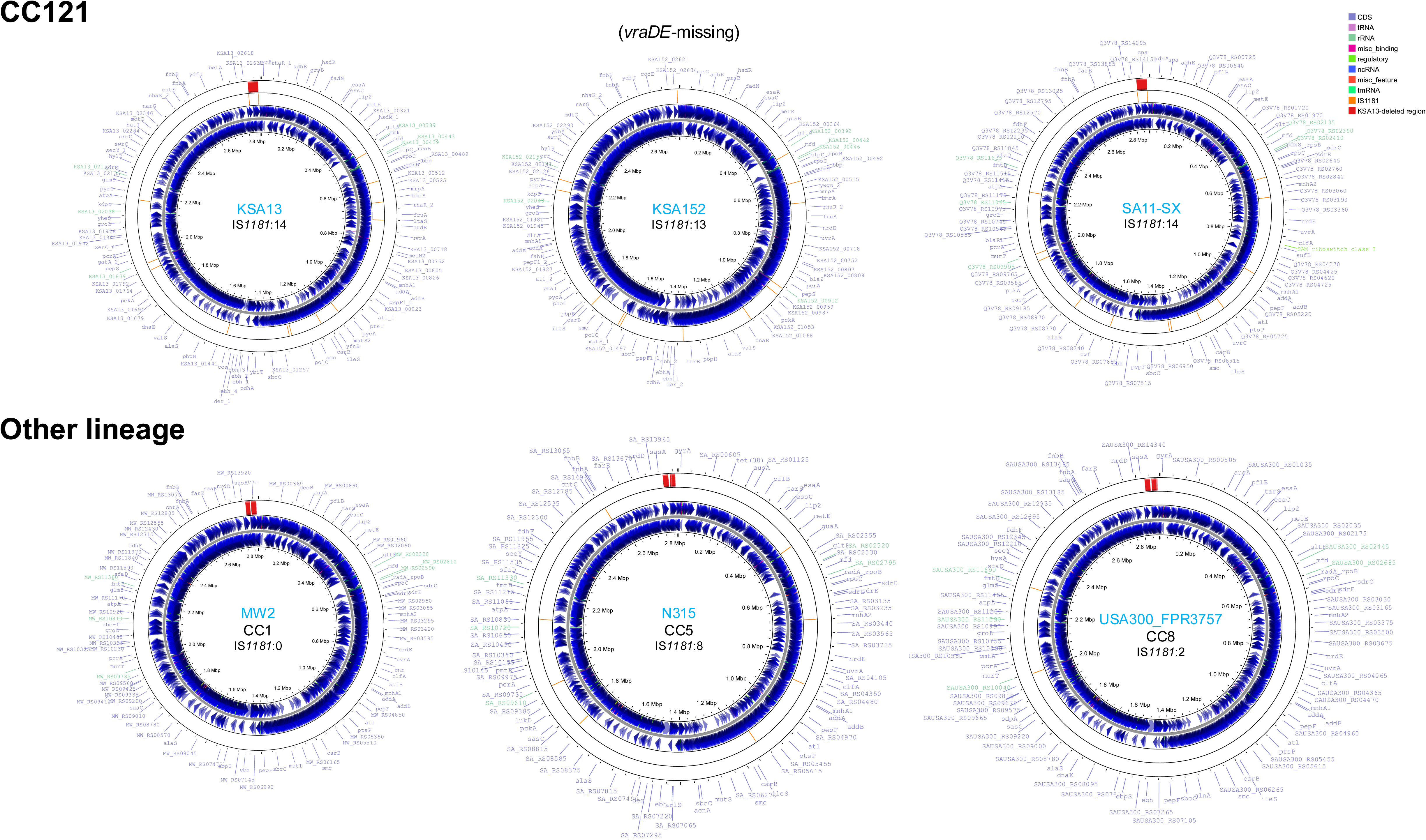
Comparison of the positions of IS*1181* and the deleted region in complete *S. aureus* genomes. The whole circles represent the chromosomes of each *S. aureus* strain. The blue arrows represent the gene loci. The orange line inside the second circle from the outside indicates the positions of IS*1181*, and the red box in the outermost circle represents the position of the deleted region, including the *vraDE* genes. The strain name, CC type and number of IS*1181* sequences on each chromosome are shown in the center of the circle.

## Discussion

In this study, we found that BraRS, BraAB and VraDE strongly affect the susceptibility of *S. aureus* to epidermin, and spontaneous mutations or deletions of these genes increase the susceptibility of *S. aureus* clinical isolates to epidermin. Epidermin is a class I type A bacteriocin that binds to lipid II and inhibits cell wall synthesis (28). This peptide was first found in *S. epidermidis* strain Tü3298 in 1986 (29), and several studies investigating the efficacy of epidermin and its derivative gallidermin were performed (30, 31). We also previously isolated an epidermin-producing *S. epidermidis* strain, KSE56, and determined the complete genome sequence of the epidermin-encoding plasmid (32). Previous studies have demonstrated that epidermin has antimicrobial activity against more than 80% of the tested *S. aureus* strains (7, 8), but 77% of our *S. aureus* strains did not show any inhibitory zone in a direct assay using KSE56 (Fig. 1). Since the amino acid sequence of epidermin in KSE56 was completely the same as that in Tü3298 (32), the difference is possibly due to the difference in epidermin-producing ability between Tü3298 and KSE56.

BraRS, a TCS that mainly upregulates the efflux transporter *vraDE*, is known to confer resistance to some antibiotics and bacitracin (9, 11). This upregulation requires both functional BraRS and BraAB (12). In the epidermin-susceptible strains, the induction of *vraD* expression in the presence of nisin A was lost or reduced. The higher susceptibility to epidermin was, the less induction of *vraD* was observed, while relatively low-susceptibility strains and MW2 tended to induce *vraD* expression (Fig. 3), suggesting that the dysfunction of BraRS systems occurred in strains highly susceptible to epidermin. Analysis of the amino acid sequences of BraRS and BraAB revealed several highly susceptible strains with frameshifts and nonsense mutations in *braR, braA* and *braB* (Fig. 4). These mutations are predicted to significantly alter the subsequent amino acid sequence or terminate the translation into amino acids prematurely, potentially causing significant changes in protein conformation or incomplete protein synthesis. However, some strains were highly susceptible to epidermin and loss of *vraD* upregulation without frameshifts and nonsense mutations. In some strains, strain-specific amino acid substitutions were detected in BraS, BraA, and BraB (Fig. 4), and these substitutions may cause dysfunction of the BraRS system. In addition, substitutions common to several strains were also observed. To investigate whether such substitutions affect the susceptibility to epidermin, we constructed a phylogenetic tree of the amino acid sequences of strains without frameshifts, nonsense mutations and *vraDE* deletions (131 strains) and evaluated the correlation with susceptibility (Fig. S5). One clade of BraR (lower part of the tree) had a relatively high proportion of susceptible strains, but no obvious correlation between susceptibility and the phylogenetic tree was observed. Therefore, the cause of impaired BraRS function in epidermin-susceptible strains has not been fully elucidated.

Deletion of *vraDE* genes also resulted in increased susceptibility to epidermin. This deletion was specifically found in CC121 strains, and the biofilm-related *ica* operon and histidine biosynthesis *his*-operon were also deleted (Fig. 5). In the deletion strains, histidine auxotrophy was observed in CDM (Fig. 6), but the strength of biofilm formation did not differ (Fig. S3). In *S. aureus*, *ica*-independent biofilm formation mediated by extracellular proteins has been reported (33, 34). These mechanisms may contribute to the biofilm formation of *ica-*negative strains. Subculture of KSA13 in 10 passages resulted in the deletion of the *ica*-operon, *his*-operon and *vraDE* genes in some colonies, suggesting that this deletion is likely caused by recombination among the two copies of IS*1181* (Fig. 5, 7C). A similar phenomenon has been reported in the ST398 MRSA strain, in which the loss of the *mecA* gene and its conversion to MSSA were observed as a result of the recombination between *ccrC* genes within the SCC*mec* cassette (35). However, to the best of our knowledge, this study is the first report of the loss of core genes due to recombination among identical IS sequences in *S. aureus*. Multiple copies of IS*1181* are known to be present on *S. aureus* chromosomes (26), but IS*1181* sequences that caused *vraDE* deletion were observed only in CC121 strains (Fig. 8). The fact that the *vraDE-*lacking strains were found only in CC121 may be attributed to this specificity. The deletion of multiple genes through recombination has the potential to result in the loss of essential genes for bacterial growth, which may be disadvantageous to bacterial survival; however, in this study, the same genomic deletion was confirmed in three strains from different sources. Thus, it seems that at least these strains can colonize the nasal cavity or cause bloodstream infections.

There are several limitations in this study. First, it was difficult to detect strains whose susceptibility to epidermin was particularly low. To gain a deeper understanding of the diversity of epidermin susceptibility in clinical isolates, experiments that are able to detect the differences in epidermin susceptibility in strains determined to have no halo in this study are needed. Furthermore, despite the findings of *vraDE-*deleted strains in our collection, no such strains were found in the 1406 complete genomes of publicly available *S. aureus*. This difference may be caused by sampling bias in strains with complete genomes because the determination of complete genome sequences is sometimes performed against high-virulence or antimicrobial-resistant strains. Consistent with this prediction, the percentage of MRSA (*mecA* gene-positive strains) was 52.7% in the complete genome collection, while the prevalence of MRSA carriage in the Asia-Pacific region ranged from approximately 0% to 40% (36), and that of our collection was 19.0%, suggesting potential bias in the collection of complete genomes. In addition, geographical and temporal bias may influence the results because our collection is limited to strains isolated in Japan in the 2010s and 2020. Further study is needed to elucidate the exact prevalence of the *vraDE-*deleted strain.

In conclusion, we investigated the diversity of epidermin susceptibility in *S. aureus* clinical isolates and elucidated the underlying mechanisms. In some strains with particularly high susceptibility to epidermin, we found that the BraRS gene and its regulon gene contained lethal mutations or that the *vraDE* gene was missing because of recombination between IS*1181*. We expect that these findings may contribute to understanding the genomic mechanisms involved in the diversity of susceptibility to antimicrobial substances in *S. aureus* populations.

## Materials and Methods

### Bacterial strains and growth conditions

The bacterial strains used in this study are shown in Table S1. KSA and HSA strains were isolated from the nasal cavity of healthy volunteers and outpatients, respectively, in the dental department (19, 20). Another 53 strains were isolated from patients with bloodstream infections (21). All the strains were isolated from independent people. *Staphylococcus* strains were grown in trypticase soy broth (TSB) (Becton, Dickinson and Company [BD], Franklin Lakes, NJ, USA) at 37 °C with shaking. When necessary, 5 µg/mL tetracycline (Wako Pure Chemical Corporation, Osaka, Japan) was added to the TSB for the *S. aureus* gene-inactivated mutants.

### Direct assay

To evaluate the susceptibility of *S. aureus* strains to bacteriocin, we performed a direct assay as described previously (11). *Staphylococcus epidermidis* KSE56 was used as an epidermin producer. The exact data were calculated from the mean diameter of the inhibition zone (halo) in three independent experiments.

### Determination of whole-genome sequences of *S. aureus* strains

DNA extraction was performed to investigate the whole-genome sequence of the *S. aureus* strains. This procedure was almost the same as that previously described (20). *S. aureus* cells grown overnight in 5 ml of TSB were collected by centrifugation and suspended in 0.5 ml of CS buffer (100 mM Tris-HCl [pH 7.5], 150 mM NaCl, and 10 mM EDTA) containing lysostaphin (Sigma‒Aldrich, St. Louis, MO, USA) (final concentration: 70 μg/ml). After incubation at 37 °C for 2 h, 0.5 ml of saturated phenol was added, and the samples were mixed well. Afterwards, the supernatant was collected by centrifugation. An aliquot of 0.5 ml of phenol‒chloroform was added to the supernatant, the sample was mixed well, and the supernatant was again collected after centrifugation. DNA was precipitated by adding an equal volume of ethanol. After centrifugation, the precipitates were dissolved in distilled water. DNA sequencing of all the *S. aureus* strains was performed with an Illumina HiSeq X FIVE. To determine the complete genome sequence, the genomic DNA of KSA13, KSA152, HSA44 and 33009-19-S058 was extracted using lysostaphin and a nucleobond HMW DNA extraction kit (Macherey-Nagel, Germany) according to the manufacturer’s protocol. DNA sequencing using PacBio HiFi reads was performed at Macrogen Japan Corporation (Tokyo, Japan), and the resulting fastq files were assembled with Illumina reads using the hybracter v0.11.2 (37).

### Analysis of genomic information

On the basis of the whole-genome sequences, we performed several genomic analyses. Multilocus sequence types (MLSTs) and clonal complexes (CCs) were determined using PubMLST (https://pubmlst.org/). Comparisons of DNA sequences among *S. aureus* strains were carried out using nucleotide BLAST (BLASTn) v2.15.0+ (38) to obtain specific sequences of targeted genes and MAFFT v7.520 (39) to align them. Maximum likelihood phylogenetic trees were constructed using RAxML-NG v1.2.0 (40) with a model LG based on the aligned amino acid sequences: BraR, BraS, BraA and BraB. The annotation of the genomes was performed using Prokka v1.14.6 (41). Genetic comparisons were performed using Easyfig v2.2.5 (42). Protein domain analysis was performed using a SMART domain search (http://smart.embl-heidelberg.de/). The complete genome sequences of *S. aureus* available in the NCBI database were obtained using the ncbi-genome-download tool from the NCBI Refseq FTP server with an assembly level ‘complete’ in FASTA format. To visualize the position of IS*1181* and the deleted region, we used the Proksee online tool (43). The genomic information of *S. aureus* MW2 (NC_003923.1), N315 (NC_002745.2), USA300_FPR3757 (NC_007793.1), and SA11-SX (NZ_CP130538.1) was obtained via Proksee.

### Purification of epidermin

Epidermin purification was performed as described previously (32). In brief, a cation-exchange resin was added to the supernatant of an overnight culture of KSE56 and stirred for one night. Subsequently, epidermin was eluted from the resin using acetic acid. The eluted samples were evaporated and dissolved in distilled water and subjected to high-performance liquid chromatography (HPLC) with an octadecyl C18 column. The fractions were collected, and their antimicrobial activity and peptide concentration were determined.

### Quantitative PCR

Quantitative PCR was performed to evaluate gene expression. When the OD_660_ reached 0.6, epidermin or nisin A (128 µg/mL) was added to the culture. Nisin A was purchased from Sigma‒Aldrich. After incubation for 30 min, the bacterial cells were collected by centrifugation. RNA extraction, cDNA synthesis, and quantitative PCR were performed as described previously (11). Gene expression levels were calculated as the ratio of total *gyrB* expression. The primers used in this assay are listed in Table S2. cDNA samples were obtained from three independent samples, and the mean value was determined as the exact gene expression level.

### Histidine auxotrophy

We used chemically defined medium (CDM) with or without histidine for the growth medium to evaluate the histidine auxotrophy of *S. aureus* strains. The method for preparing CDM was described previously (44). The overnight culture of the *S. aureus* strains was adjusted to an OD_660_ of 1.0, and the cells were collected by centrifugation. After the cells were washed with phosphate-buffered saline (PBS), 1 × 10^8^ cells were added to 3 mL of CDM with or without 1 mM L-histidine (Katayama Kagaku Industries, Osaka, Japan) and incubated at 37 °C for 24 h with shaking.

### Quantification of biofilm formation

To quantify the biofilm formation of *S. aureus* strains, we performed a crystal violet staining assay as described previously with minor modifications (19). Aliquots of 100 μl of TSB containing 10% glucose were added to a 96-well flat bottom plate, and 1 × 10^5^ *S. aureus* cells were inoculated overnight and incubated for 24 h at 37°C. The biofilms were washed with PBS three times and subsequently stained with 0.1% crystal violet solution for 5 min. After rinsing with PBS, the cells were eluted with 30% acetic acid solution, and the absorbance (A_570_) of the solution diluted 5-fold with water was determined using an iMark Microplate reader (Bio-Rad Laboratories, CA, USA).

### Isolation and analysis of the *ica, his, vraDE*-deleted mutant of KSA13

To isolate the mutant that requires histidine for growth from KSA13 culture, CDM agar plates with or without 1 mM histidine were prepared. After KSA13 was cultured through 10 passages in TSB with 1 mM histidine, the culture was appropriately diluted and plated on trypticase soy agar (TSA) and incubated for 16 h. The colonies that appeared were replated on CDM plates and incubated for 48 h. The colonies that grew in CDM with histidine but not in CDM without histidine were selected, and the deletions of *icaA, hisD* and *vraD* in the mutants were confirmed by PCR using specific primers (Table S2). To sequence the missing region, the primers that annealed upstream of *icaR* and downstream of *cspB* were used for PCR (Table S2), and the obtained amplicon was subsequently purified using a QIAquick PCR purification kit (QIAGEN, Tokyo, Japan). Sangar sequencing was performed at Macrogen Japan Co. using 8 primers (Table S2), and the resulting sequences were integrated.

### Statistical analysis

Dunnett’s multiple comparison test was performed to compare the halo sizes shown in Fig. 2. Šidák’s multiple comparison test was performed to compare *vraD* expression, as shown in Fig. 3. Tukey’s multiple comparison test was performed to compare *vraD* expression, as shown in Fig. S1. Student’s *t* test was performed to compare A_570_, as shown in Fig. S3. All the statistical analyses were performed with GraphPad Prism (GraphPad Software, San Diego, CA, USA).

## Abbreviations

MRSA: methicillin-resistant *Staphylococcus aureus*
MSSA: methicillin-sensitive *Staphylococcus aureus*
CC: clonal complex
CDM: chemically defined medium

## Data availability

The complete genomes of the KSA13, KSA152, HSA44 and 33009-19-S058 strains have been deposited in the DNA Data Bank of Japan (DDBJ) under bioproject number PRJDB35750. The whole-genome sequences of *S. aureus* strains in this study have been deposited in DDBJ database under following bioproject number; KSA strains: PRJDB15626; HSA strains: PRJDB18935; Strains from bloodstream infection: PRJDB15501.

## Acknowledgement

We declare that there are no conflicts of interest.

This study was supported in part by Grant in-Aid for Scientific Research (C) (Grant No. 24K12886) from the Ministry of Education, Culture, Sports, Sciences and Technology of Japan. This study was also supported in part by JST SPRING, Grant Number JPMJSP2132.

## CRediT Author Contributions

Conceptualization: Y.S., M.K.-M. and H.K. Data curation: Y.S., A.O. and H.K. Formal analysis: Y.S. and A.O. Funding acquisition: Y.S., M.K.-M. and H.K. Investigation: Y.S., A.O., J.H., C.A., S.N. and H.K. Methodology: Y.S., J.H. and H.K. Project Administration: Y.S., M.K.-M. and H.K. Resources: Y.S., M.K.-M., J.H., C.A., M.S. and H.K. Software: Y.S. Supervision: M.S., S.Y., T.S. and H.K. Validation: Y.S., A.O., and H.K. Visualization: Y.S. and A.O. Writing-original draft: Y.S. and A.O. Writing-review and editing: All authors read, sub-edited, and approved this manuscript.

## Supplemental materials

Table S1. Bacterial strains used in this study.

Table S2. Primers used in this study.

Fig. S1 Gene expression of *vraD* with 128 μg/mL nisin A or epidermin in WT MW2 and the Δ*braRS* mutant.

nis+: 128 μg/mL nisin A; epi+: 128 μg/mL epidermin. Statistical analysis was performed using Tukey’s multiple comparison test. ****, *P* < 0.0001; ns, not significant.

Fig. S2 Proportion of each CC among the *S. aureus* clinical isolates used in this study. The numbers below each CC indicate the number of strains belonging to that CC.

Fig. S3 Comparison of biofilm formation among CC121 strains.

*icaRADBC +* includes the 6 strains with *ica* genes, and *icaRADBC* – includes the 3 strains without *ica* genes. Biofilm formation was quantified by measuring the absorbance at 570 nm. The exact absorbance was calculated from three independent experiments. Statistical analysis was performed using Student’s *t* test. ns, not significant.

Fig. S4 Numbers of IS*1181* copies in strains of each representative CC.

The black points show the number of IS*1181* copies in complete *S. aureus* genome sequences obtained from the NCBI Refseq database. The number of IS*1181* copies was determined using NCBI BLASTn.

Fig. S5 Phylogenetic trees of the amino acid sequences of BraR (A), BraS (B), BraA (C) and BraB (D) and heatmaps of the halo size of epidermin in 131 *S. aureus* clinical strains without frameshifts, nonsense mutations and deletion of *vraDE*.

